# MS-BCR-DB: an integrated BCR repertoire database to mine humoral multiple sclerosis signatures

**DOI:** 10.64898/2026.03.05.709606

**Authors:** Chiara Ballerini, Niccolò Cardente, Maria Francesca Abbate, Khang Lê Quý, Natalia Rincon, Lina Wolfram, Andreas Lossius, Emilio Portaccio, Maria Pia Amato, Clara Ballerini, Victor Greiff

## Abstract

Multiple sclerosis (MS) is a chronic autoimmune disease of the central nervous system (CNS) in which B cells play a critical role. While B-cell receptor (BCR) sequencing studies in MS are increasing, progress in understanding MS-associated BCR repertoire features and convergent patterns across patients has been limited by small cohorts, heterogeneous experimental methodologies, and fragmented data storage. To overcome these challenges, we developed the MS-BCR-Database, the first publicly accessible and uniformly processed collection of human MS BCR sequencing datasets. We harmonized raw BCR-sequencing data into an AIRR-compliant database incorporating clinical and technical metadata, enabling coherent cross-study analyses. Using this resource, we identified putative disease-associated BCR-sequence features, including CNS-biased V-gene usage, marked oligoclonal expansion in cerebrospinal fluid, and convergent clonotype clusters shared exclusively among MS patients. Integration with antigen-annotated BCR databases revealed matches to antibodies recognizing both viral antigens, including Epstein-Barr virus, and CNS self-proteins. The MS-BCR-Database provides a scalable foundation for mechanistic discovery and biomarker development in MS, while establishing a broadly applicable resource for integrative analyses of BCR repertoires.

## Introduction

Multiple sclerosis (MS) is a chronic demyelinating autoimmune disease of the central nervous system (CNS) that usually affects young adults with a median age of onset around 30 years. The disease is clinically heterogeneous and encompasses distinct disease courses, including clinically isolated syndrome (CIS), radiologically isolated syndrome (RIS), relapsing-remitting MS (RRMS), primary progressive MS (PPMS) and secondary progressive MS (SPMS) (Oh et al. 2018). MS is mediated by both innate and adaptive immunity, with adaptive immune cells playing the dominant role in initiating and sustaining CNS inflammation (Afzali and Korn 2025; Dendrou et al. 2015).

The etiopathogenesis of the disease and the antigenic targets of the immune response remain unknown. While MS has been predominantly considered a T-cell mediated disease, in the last decade it has become evident that B cells are also involved, with converging evidence emerging from several independent studies, namely: the efficacy of anti-CD20 drugs in treating the disease, targeting specifically the B cells (Hauser 2015); the presence of oligoclonal bands (OCB) in the cerebrospinal fluid (CSF) of MS patients, a sign of intrathecal antibody production (Di Sabatino et al. 2025); the in vitro demonstration that B cell secretions from RRMS patients contain toxic factors for oligodendrocytes (Lisak et al. 2012) and that MS CSF-derived antibodies induce demyelination in spinal cord slices (Lisak et al. 2017); the presence of B cells in white and grey matter lesions and ectopic meningeal follicles, along with IgG and complement deposition (Magliozzi et al. 2007; Serafini et al. 2004).

B cells contribute to CNS damage in MS through several mechanisms, including antibody production, pro-inflammatory cytokine release, and acting as antigen-presenting cells (APCs) to CD4+ T cells (Hauser 2015). Their pathogenic potential has been further underscored by recent findings showing that clonally expanded B cells in MS recognize antigens of Epstein-Barr virus (EBV), a known risk factor for MS (Bjornevik et al. 2022), and cross-react with CNS proteins, providing a possible molecular link between infection and autoimmunity (Thomas et al. 2023; Marti et al. 2024; Lanz et al. 2022; Younis et al. 2026). A better characterization of these cells may therefore help elucidate the mechanisms underlying MS development.

Indeed, interest in studying the receptors responsible for the adaptive immune response has grown considerably over the last decade, particularly in the context of lymphoid and autoimmune diseases (Liu et al. 2021; Ishigaki et al. 2022). AIRR-seq represents a powerful approach to this end, enabling deep profiling of B cell receptor (BCR) and T cell receptor (TCR) repertoires with unprecedented resolution and throughput in health and disease (Mhanna et al. 2024; Greiff et al. 2015).

To date, there exists no integrated resource for BCR studies in MS. Among the studies available, some are relatively dated, relying on early sequencing technologies such as 454 sequencing (Slatko et al. 2018), or are focused on small cohorts, restricted tissue sampling, or therapy-specific cohorts (Palanichamy et al. 2014; Greenfield et al. 2019; Stern et al. 2014; Lomakin et al. 2022), providing a fragmented view of the BCR landscape in MS. In other fields, disease-specific databases such as ERIC CLL-DB (Chronic Lymphocytic Leukemia Database) and CovAbDab (COVID-19 Antibody Database) have accelerated understanding of disease and BCR characterization, while facilitating diagnostic and therapeutic advances (Chatzidimitriou et al. 2020; Raybould et al. 2021). These resources exemplify how curated databases can drive progress in diseases driven by adaptive immunity.

To address the lack of an integrated MS-specific BCR data resource, we developed a centralized database that aggregates publicly available MS BCR raw data and processes them through a uniform pipeline to harmonize results across studies. This approach expands patient representation and provides the research community with a coherent, interpretable, and harmonized resource. Although technical variations, such as sequencing platforms and library preparation protocols, are unavoidable, we standardized downstream analyses and annotations as much as possible. Leveraging this resource, we performed analyses to identify potential disease-related BCR signatures in MS patients. The MS-BCR-Database provides a foundation for future integrative studies of the adaptive immune response in MS.

## Results

### Building a harmonized MS-BCR-Database

We developed a harmonized MS-BCR-Database, integrating publicly available data into a standardized framework to enable cross-study comparisons, downstream analyses, and promote broader biological insight. From our systematic review, we identified 11 studies (Agrafiotis et al. 2023; Greenfield et al. 2019; Laurent et al. 2023; Lomakin et al. 2022; Palanichamy et al. 2014; Saldivar et al. 2022; Ramesh et al. 2020; Ruschil et al. 2023; Zvyagin, n.d.; Ryback and Cowan 2025; Stern et al. 2014) (flowchart Fig. 1A). Inclusion criteria included human MS BCR studies with accessible raw sequences published after 2005. Studies on animal models, non-MS cohorts, or lacking raw data were excluded.

**Fig. 1.**
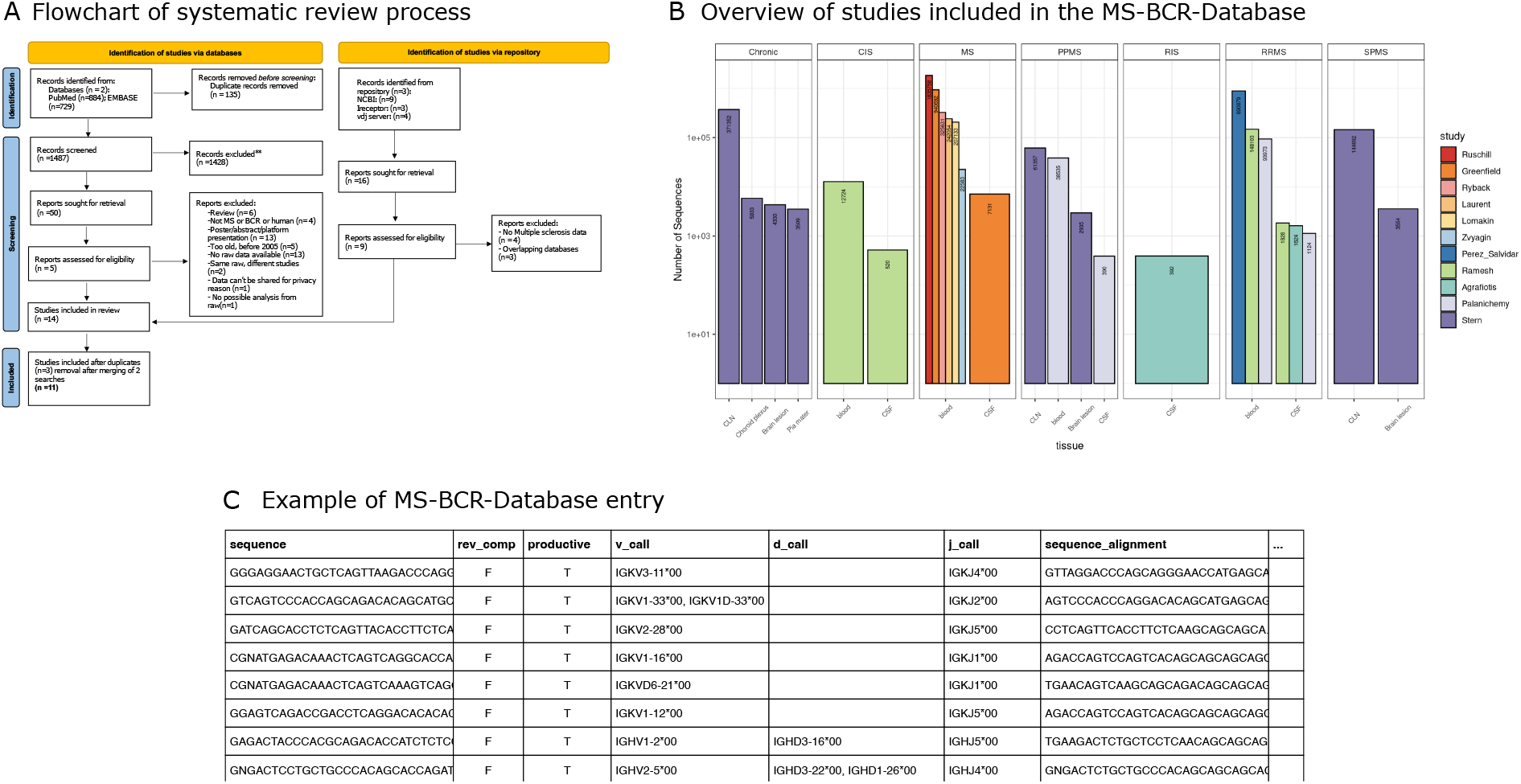
**A Flowchart of the systematic review process, adapted from the official PRISMA guidelines** (Page et al. 2021). The flow chart shows the total number of research articles screened, the databases queried, and the inclusion and exclusion criteria considered to select the final studies for the construction of a BCR-based MS-specific database. **B Overview of studies included in the database.** Bar plot showing the contribution of each study to the final database. Bars are colored by study and faceted by different MS types. The x-axis indicates tissues analyzed, while the y-axis shows the number of sequences in log scale. Numbers within bars represent the counts of sequences with the corresponding characteristics. **C Example of dataset entry**. Abbreviations CIS: Clinically isolated syndrome; MS: multiple sclerosis; RRMS: relapsing remitting MS; PPMS; primary progressive MS; SPMS: secondary progressive MS; RIS: radiologically isolated syndrome; CLN: cervical lymph nodes; CSF: cerebrospinal fluid

The 11 included studies comprised a total of 5,368,636 BCR sequences, with a mean of 47,093 sequences per patient (SD: 72,494). This variability likely reflects a highly skewed distribution of sequence counts per patient, partly driven by the diversity of tissues analyzed. The included studies investigated different types of MS patients (total number of patients = 114), either treatment-naïve or under therapy, at different time points and across multiple tissues, resulting in a heterogeneous collection in terms of clinical context and sample origin (Fig. 1B). In two studies, datasets (Ramesh et al. 2020; Agrafiotis et al. 2023) were generated using single-cell sequencing, whereas the remaining studies were based on bulk sequencing (five of which contained both heavy and light chains while the other four only heavy).

For each study, we computed mean sequence quality scores with FastQC (Supplementary table 3) (“Babraham Bioinformatics - FastQC A Quality Control Tool for High Throughput Sequence Data,” n.d.). When available, associated metadata were incorporated into the database (e.g., patient characteristics, treatment status, tissue of origin). A summary of all studies and their key features is provided in Supplementary Table 1.

### Insights from exploratory analysis on the BCR-MS-Database

#### Strategy for BCR analyses leveraging MS-BCR-DB

To showcase the investigation of disease-specific BCR signatures in MS using the BCR-MS-Database, we focused on therapy-naïve MS patients to minimize treatment-related variability. We also retained an autopsy-based study (Stern et al. 2014), which analyzed particularly informative tissues such as brain and lymph nodes; although treatment information was not reported, post-mortem samples were considered unlikely to reflect therapy effects. In addition, we restricted our analyses to heavy-chain sequences, as not all studies provided both heavy- and light-chain data. This subset of the MS-BCR-DB yielded 66 patients, 6 different tissues, yielding a total of 1,613,653 sequences after MiXCR processing and VDJ annotation, with a mean of 24,449 sequences per patient (SD 43,911). In addition, healthy controls (HCs) from Ghraichy et al. were included (38 subjects, age range 15.1-45.4 years; 3,178,078 sequences; mean sequences per patient 83,634, SD 57,234) (Ghraichy et al. 2020).

In our analysis, we focused on germline-gene-usage patterns, clonal expansion and shared disease-specific BCR clusters across patients and tissues (Fig. 2; Supplementary Materials Figs. 1-3). Such BCR features are potentially relevant to MS because the use of tissue-specific V genes can shed light on how B cells generate a persistent immune response within the CNS, informing “inside-out” versus “outside-in” models of disease development (Luchicchi et al. 2021). The identification of expanded B-cell populations, particularly within CNS tissues, may pinpoint the clones driving the autoimmune process (von Büdingen et al. 2012). Furthermore, BCR clusters shared among patients and tissues may reflect convergent responses to common antigens (Ramesh et al. 2020; Palanichamy et al. 2014; Lindeman et al. 2022). Collectively, this pilot analysis demonstrates how the MS-BCR-DB can be used for standardized, comparative immunological investigations in MS.

**Fig. 2.**
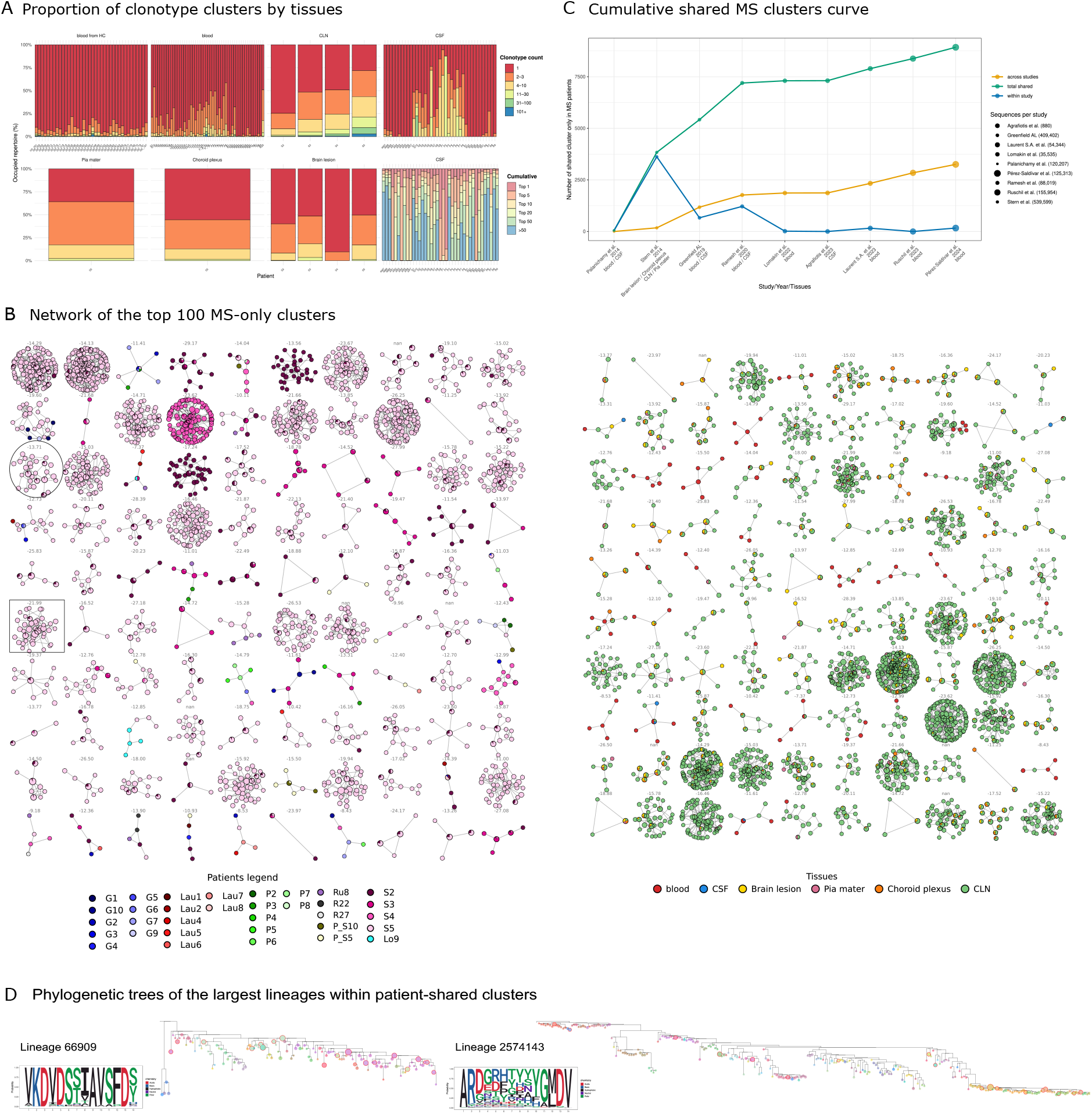
B-cell response in MS: oligoclonal CSF profile, sharing of clusters across patients, and extensive intra-patient lineage expansion. **A Proportion of cluster of clonotypes (defined as a combination of V and J genes and HCDR3 amino acid sequence) by tissue.** For each sample in different tissues, stacked bars show the fraction of the repertoire contributed by clonotypes grouped by count-based bins (1, 2-3, 4-10, 11-30, 31-100, 101+). Legend colors (right, “Clonotype count”) map to these bins (e.g., 1 = singleton, red). Bottom-right, in lighter tones, the CSF panel is shown in a rank-based view, indicating how much of the repertoire is explained by the most abundant clonotypes versus the remainder (>50). In CSF, low-abundance clonotypes contribute minimally, indicating that most of the repertoire is dominated by a few highly expanded clonotypes (oligoclonal profile). **B Network of the top 100 largest clusters shared by at least two different MS patients**. Clusters seen in healthy controls were removed, so everything shown is MS-only. Each node represents a unique V-HCDR3-J amino-acid sequence; edges link sequences that differ by one amino acid (Hamming distance = 1). Node size increases with the number of observations of that sequence in the filtered data (log-scaled). In the left plot, nodes are colored by subject; on the right, nodes are colored by tissue. When the same sequence is observed in multiple subjects (or multiple tissues), the node is drawn as a pie chart with slice sizes proportional to their counts. **C Cumulative shared MS clusters curve**. Studies are ordered along the x-axis by year of publication. The y-axis reports the number of clusters shared by at least two patients. Dot size reflects the exact number of BCR sequences analyzed in each study, considering clusters detected exclusively in MS patients. The three curves represent clusters shared within the same study, clusters shared across different studies, and the total number of shared clusters (union of within- and across-study sharing, with clusters counted uniquely). **D Reconstructed phylogenetic trees of the largest lineages within patient-shared clusters**. Phylogenetic trees for lineage IDs 66909 and 20719. Lineage 20719 is part of the circle network, while lineage 66909 belongs to the framed network in Fig. 2B. Each circle represents a unique amino-acid sequence and is colored according to its HCDR3 amino-acid composition; different sequences correspond to different colors. Node size depends on the number of nucleotide sequences. Nodes highlighted with red circles are those for which the same nucleotide sequence has been found in different tissues (all include CLN and at least one intracranial tissue). To the left of each tree, the lineage-specific HCDR3 amino-acid sequence logo shows positional variability and highlights enriched amino acids at each position. Both lineages originate from patient S5. Abbreviations HC: healthy controls; CSF: cerebrospinal fluid; CLN: cervical lymph nodes.

#### Evidence for tissue-compartmentalized BCR repertoire signatures in MS

We aimed to determine whether specific features of BCR in MS could identify distinct patterns of immune activation and compartmentalization in different tissues, especially within the CNS. We examined the distribution of HCDR3 heavy chain length and V-D-J gene usage across different tissues (Supplementary Materials, Fig. 1-2), and compared them with HCs (Supplementary Materials, Fig. 3). An enrichment (mean 31.0% vs 21.8%; +9.2 percentage points; 1.42-fold; p_adj_ < 0.05) of the IGHV4 family was observed in intracranial samples compared to peripheral ones (Supplementary Materials, Fig. 2B). This finding is consistent with previous evidence suggesting that IGHV4 is preferentially utilized by clonally expanded B cells in MS intracranial tissue, and these cells produce antibodies that target CNS antigens, contributing to the pathogenesis of MS (Owens et al. 2007; Rivas et al. 2017). In addition, IGHJ5 usage was higher in intra-tissue B cells than in extra-tissue samples (p_adj_ < 0.01; mean 14.2% vs 11.8%; +2.4 percentage points; 1.22-fold increase).

We also observed a significant overexpression (mean 6.66% vs 4.07%; +2.59 percentage points; 1.64-fold increase; p_adj_ < 0.01) of usage of IGHV1-69, a gene linked to antiviral and autoreactive responses (Watson and Breden 2012; Al Kindi et al. 2015; Shiroishi 2021), in MS patients compared to HCs (see Supplementary Materials Fig. 3A). These findings are consistent with an enrichment of potentially autoreactive-prone B cell clones in MS (Li et al. 2018; Bashford-Rogers et al. 2018). A significant reduction in IGHV4-34 usage was detected in MS patients compared to HCs (p_adj_ < 0.001; mean 4.19% vs 8.23%; −4.04 percentage points; 0.51-fold change).

We examined the distribution of clonotype clusters across tissues for each patient using single-linkage clustering. A clonotype was defined as a unique combination of V and J genes and HCDR3 amino acid sequence. Clonotypes were grouped into the same cluster when they shared identical V and J genes and were connected by a link if they differed by one amino acid in the HCDR3 (Hamming distance =1) (Spisak et al. 2022). In CSF, highly expanded clonotype clusters dominated the repertoire. Size rank-based analysis showed that the top clonotype cluster accounted for 11.1% of all sequences, while clonotype clusters ranked 2-5 contributed an additional 20.7%; altogether, the top 10 clusters represented the 42.6% of the repertoire. In contrast, the long tail of low-abundance (ranks >50) contributed only 26.4%. Consistently, abundance-binned analysis demonstrated an overrepresentation of expanded clonotype clusters: the clusters comprising 11-30 or 31-100 sequences represented 25.2% and 22.2% of the repertoire, respectively. In peripheral blood, differences between MS patients and HCs were present but modest. Both groups displayed a predominantly polyclonal architecture, dominated by low-abundance clonotype clusters (1-3 sequences per cluster: 79.9% in MS; 90.4% in HCs) (Fig. 2A).

#### Identification of shared and convergent BCR clusters across MS patients

Enabled by access to the MS-BCR-Database, we next examined whether MS patients shared exclusive clusters of B-cell clones potentially indicative of disease-associated convergent responses. To this end, we analyzed the shared unique amino acid clones across different patients and HCs by pooling all data and applying a single-linkage clustering algorithm, in which a link connects two sequences if they share identical V and J genes and their HCDR3 regions have a Hamming distance of 1 (Spisak et al. 2022). Unlike the within-subject linkage clustering described above, this analysis was performed across all subjects. By removing clusters containing clones present in at least one healthy subject, we identified 8927 clusters that were exclusively present in MS and absent in HCs, totaling 59504 sequences. Since clusters containing sequences from HCs were excluded, the shared clusters analyzed here represent patient-specific convergent rearrangements, potentially reflecting common antigenic stimulation underlying the disease. Among these, 3689 were restricted to blood, 704 to CLN, and 3042 were shared across intra- and extracranial compartments. To focus on biologically relevant signatures of possible disease-associated convergent structures, we developed a composite scoring framework to rank clusters by their degree of sharing across patients and tissues (adjusted for distributional evenness across subjects) and cluster size (see methods), and selected the top 100 clusters for visualization (Fig. 2B). Notably, the number of clusters shared among MS patients increased progressively with the inclusion of additional studies, indicating that data aggregation substantially enhances the detection of convergent B-cell responses and increases the power to identify putative disease-associated immune signatures (Fig. 2C).

To further characterize clonal relationships, we inferred lineages for each patient using the HILARy software (see methods) (Spisak et al. 2022). The mean number of distinct lineages per patient was 19,063 (SD=26,338). The observed inter-individual variability likely reflects differences in tissue availability across patients. In HCs, the mean number of lineages was 76,808 (SD=52,856). To investigate whether similar HCDR3 sequences observed across different patients originated from comparable clonal processes or from independent recombination events, we correlated the HILARy-defined clonal lineages (computed within each subject) with the inter-patient clusters. After excluding clusters shared with HCs, we identified 18,457 subject-specific HILARy-defined clonal lineages that participated in inter-patient clusters involving at least two subjects, indicating either shared common ancestors across patients or convergent clonal processes.

We reconstructed and visualized the lineage trees of two lineages with the greatest number of branching events (Fig. 2D). These highly expanded lineages are indicative of somatic hypermutation and affinity maturation, including prolonged B-cell activation (Blazso et al. 2022; Horns et al. 2016). We observed that some identical nucleotide sequences present within the lineage are shared across multiple tissues (highlighted in red in Fig. 2D). All of them occur in both extracranial and intracranial compartments, specifically CLN and at least one of the following: brain lesions, choroid plexus, or pia mater. Both reconstructed lineages originate from clusters shared only among MS patients (specifically, the two networks highlighted by the black square and the circle in Fig. 2B), illustrating how clonotype clusters shared across different subjects can include highly expanded and evolutionarily diversified lineages within individual patients. Notably, previous work, using quantitative lineage tree analysis, has shown that autoimmune lineage trees tend to be larger than those observed in HCs (Steiman-Shimony et al. 2006).

To evaluate whether the presence of shared HCDR3 amino acid sequences across patients might reflect convergent selection events rather than stochastic recombination, we estimated the generation probability (Pgen) (Sethna et al. 2019) of all inter-patient shared clonotypes. We specifically focused on the subset of sequences shared by at least two MS patients and absent in HCs (same set of sequences mentioned above), resulting in a group of ∼30347 candidate clonotypes with low recombination likelihood (Pgen < 10^−16^). In addition, we observed a statistically significant difference in Pgen between sequences shared only among MS patients and those present in both MS patients and HCs (p < 2.2 × 10^−16^; median Pgen MS-only = 5.57 × 10^−17^; median Pgen MS and HCs = 8.41 × 10^−10^, representing a ∼10,00,000-fold difference in generation probability). Such low-probability sequences are unlikely to arise by chance, suggesting that common antigen-driven selection may underlie their emergence (see Supplementary Material Fig. 4). The complete list of these convergent, MS-associated clonotypes is provided in Supplementary Table 2 and serves as a resource for further investigation of potential disease-relevant antibody specificities.

### Mining antigen-specific sequences in MS-BCR-DB

To determine whether MS-associated BCR clones recognize known antigenic targets, we focused on HCDR3 amino acid sequences uniquely found in MS patients and absent in HCs, and compared them with the AgAb database (https://naturalantibody.com/agab/) (Czerwiński et al. 2025) of antigen-specific antibodies. A total of 186 MS-derived HCDR3 sequences showed statistically significant similarity to entries in the AgAb database (p_adj_ < 0.05), including 3 exact matches (Levenshtein distance = 0), 59 near-identical matches (distance = 1), and 124 similar matches (distance = 2) (Fig. 3). Among the matched antibodies, we identified both exogenous and endogenous targets. Exogenous antigens included viral, bacterial, toxin-associated, and parasitic proteins, whereas endogenous antigens included those from both the central nervous system and peripheral tissues. Among endogenous CNS-related antigens, MS-associated HCDR3 sequences showed matches to antibodies targeting proteins implicated in axonal growth regulation, synaptic signaling, blood-brain barrier dynamics, and neuron-glia interactions. These included Reticulon-4 (Nogo-A) and its receptor, key inhibitors of axonal regeneration following CNS injury (Kurowska et al. 2014; Rashidbenam et al. 2023; Satoh et al. 2005) as well as Neurogenic locus notch homolog protein 1 (NOTCH1), a central regulator of neurodevelopment and glial activation (Mora and Chapouly 2023; John et al. 2002). In addition, matches were observed against the Glutamate receptor ionotropic NMDA subunit 1, a core component of excitatory synaptic transmission implicated in neurodegeneration and synaptic dysfunction (Centonze et al. 2010; Rossi et al. 2013) and Annexin A1, a key mediator of inflammation resolution and blood-brain barrier integrity (McArthur et al. 2016). Among viral antigens, we identified MS-associated HCDR3 sequences matching antibodies recognizing Epstein-Barr virus proteins (EBNA1).

**Fig. 3.**
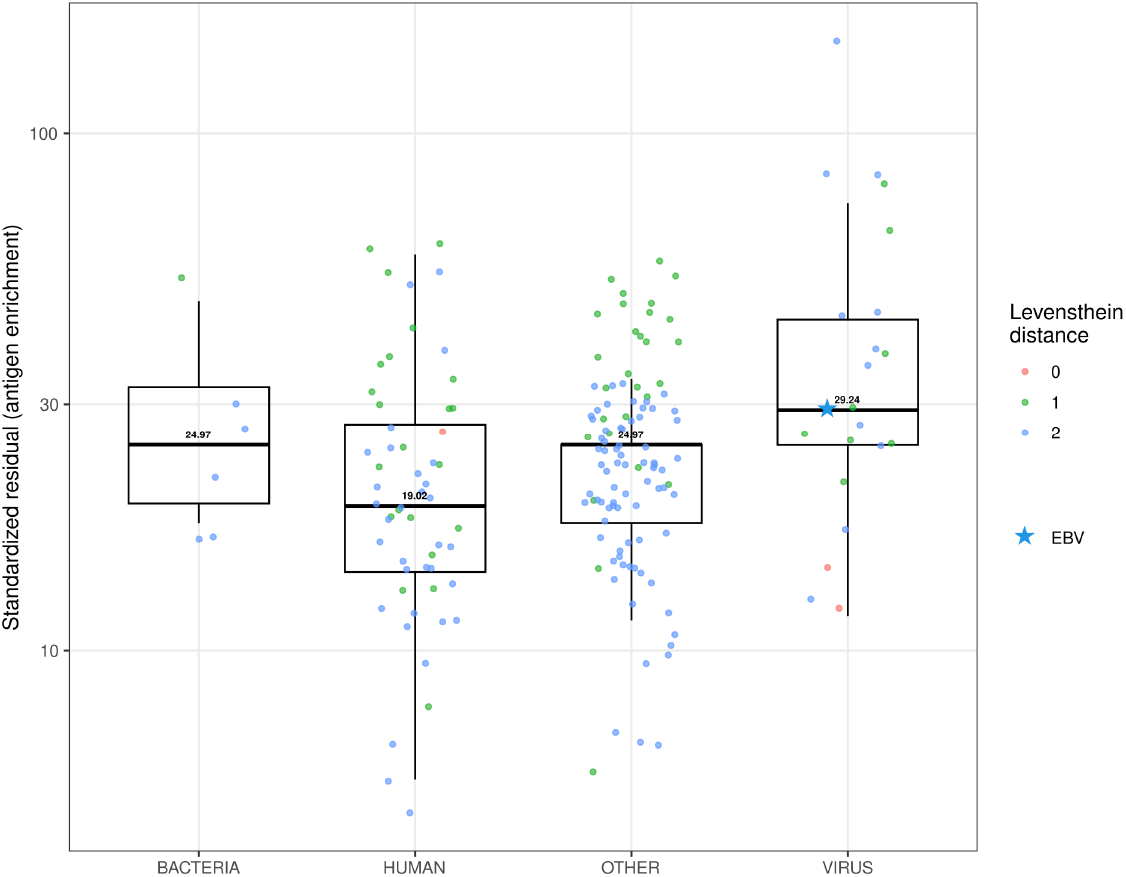
Integration of MS-BCR-DB with the antigen-specific database AgAb. Standardized residuals from Fisher’s exact tests were used to quantify the degree of over-representation of matched MS sequences relative to random expectation for each sequence with antigen annotation in the AgAbDB (Czerwiński et al. 2025; Amoriello et al. 2021). Only matches with MS-exclusive clusters and significant enrichment (FDR-adjusted P < 0.05) are shown. Positive standardized residuals on y-axes (log scale) indicate antigen-specific enrichment in the repertoire. The facet depends on the difference of Levenshtein distance (0-1-2). Each box summarizes the distribution of enrichment values for all antigens within a category, while individual points represent a single antigen sequence. In total, 55 human, 22 viral, 6 bacterial, and 103 “other” antigen-associated sequences were identified within the significant MS-restricted set. The “other” category includes antigens of parasitic, murine, primate (non-human), and additional heterogeneous taxonomic origins. The starred point indicates a match to the EBV antigen EBNA1.

These findings support the possible presence of antigen-driven B-cell responses in MS, involving both self-reactivity and cross-reactivity to infectious agents.

## Discussion

To our knowledge, the MS-BCR-DB is the first publicly available, uniformly processed MS BCR database that integrates all data currently MS-related BCR sequencing data accessible in the literature.

The strength of the MS-BCR-Database lies not only in its size but also in its comprehensiveness, encompassing samples from diverse tissues, disease phenotypes, and patient demographics. Furthermore, all datasets were analyzed from scratch using a uniform, harmonized pipeline, ensuring reproducibility and enabling fair comparisons across studies. As demonstrated across biomedical research, large and heterogeneous datasets enhance reproducibility, facilitate model validation, and foster new discoveries (Guo 2015; Cremin et al. 2022). Disease-specific repositories, such as the ERIC CLL database and Cov-AbDab, have already demonstrated the translational impact of such integrative efforts (Raybould et al. 2021; Chatzidimitriou et al. 2020). To ensure data quality and interoperability (Peng et al. 2022), we followed AIRR Community recommendations, achieving full FAIR (Findable, Accessible, Interoperable, Reusable) compliance. We provide a per-dataset sequence quality score and straightforward online download access to our ready-to-use database, offering users a standardized indicator of sequencing quality to support transparency and broad use.

We used the MS-BCR-DB to investigate alterations in the BCR repertoire in MS and identify shared immunological signatures. We observed a significant overrepresentation of the IGHV4 family in intracranial compared to peripheral tissue, in line with previous reports (Baranzini et al. 1999; Owens et al. 2006). The CSF repertoire shows extensive clonal expansion, in contrast to the broader, more polyclonal peripheral repertoire (Qin et al. 1998; Bankoti et al. 2014). The persistence of IGHV4-biased, clonally related B cells in CNS compartments supports the concept of a compartmentalized, antigen-driven immune response. Although the functional relevance of this IGHV4 bias remains unclear, IGVH4^+^ plasmablasts from MS patients have been shown to target neuronal and astrocytic antigens, suggesting recognition of brain-specific epitopes. This may reflect either early disease triggers or mechanisms sustaining chronic CNS inflammation (Rivas et al. 2017).

We also found a statistically significant increase in the use of the IGHV1-69 gene, alongside a reduction in the use of the IGHV4-34 gene, compared with HCs. Although increased IGHV4-34 usage has also been reported in some MS cohorts (Jelcic et al. 2025), the literature is not entirely consistent. Other studies have instead observed a significant decrease in IGHV4-34 usage in MS compared with both HCs and patients with other inflammatory neurological diseases (OIND), whereas in OIND an increase was observed compared with HCs (Saldivar et al. 2022). IGHV1-69, although not typically dominant in MS, is widely used in antiviral responses and has been linked to polyreactivity and autoreactivity in inflammatory and autoimmune diseases (Avnir et al. 2016; Watson and Breden 2012). Antibodies encoded by this gene frequently target conserved viral epitopes and exhibit cross-neutralizing activity, mediated by key anchor residues in the CDR-H2 and CDR-H3 regions (Avnir et al. 2014). Notably, genetic variation affecting IGHV1-69 regulation includes a promoter polymorphism overlapping a binding site for transcription factor RUNX3, whose expression is induced upon EBV infection (Avnir et al. 2016; Spender et al. 2002). In this context, it is noteworthy that MS has been strongly associated with prior EBV infection (Bjornevik et al. 2022). In the setting of chronic antigenic stimulation, the coexistence of broad viral reactivity and potentially intrinsic polyreactivity could contribute to inflammatory or autoreactive tendencies, consistent with the functional plasticity of B cells at the interface between antiviral immunity and autoimmunity. We additionally detected higher IGHJ5 expression in intracranial samples, though this segment has not been directly implicated in autoimmunity. IGHJ5 is considered a gene of medium usage in physiological repertoires (≈ 10-20 % of rearrangements) (Shi et al. 2020).

Extending our analysis, we next explored whether convergent B cell signatures are detectable across multiple MS patients. In our study, we identified approximately 8,000 BCR clonotype clusters shared exclusively among MS patients, supporting and expanding previous evidence of stereotypical or convergent B cell responses in MS (Pérez-Saldívar et al. 2024; Ramesh et al. 2020; Singh et al. 2013). The sharing of these specific clusters, absent in HCs, suggests a disease-specific convergence of the B-cell response, consistent with recurrent IGHV4 family usage, biased V-J pairings, and clonal connectivity across CSF, meninges, and parenchyma (Lovato et al. 2011; Lindeman et al. 2022; Bankoti et al. 2014). The detection of “shared-only” clusters supports the hypothesis of common antigenic or structural epitopes driving B cell activation in MS, although their precise targets remain unidentified (Singh et al. 2013; Lindeman et al. 2022).

Importantly, lineage-tree reconstruction of the most expanded MS-exclusive lineages revealed extensive branching and pronounced somatic hypermutation, features indicative of sustained antigen-driven maturation. These expanded and deeply branched trees highlight the possible biological relevance of the shared clusters and further suggest that the convergent signatures we observe could reflect genuine, disease-associated immune processes rather than stochastic overlap. Collectively, these findings motivate future antigen-mapping and functional studies to elucidate the contribution of these clusters to MS pathogenesis.

In this context, Pgen analysis provides insight into the selective forces shaping shared BCR repertoires in MS. Sequences detected exclusively in MS patients, compared with those shared with HCs, showed lower Pgen values, suggesting selective enrichment of rarer rearrangements rather than stochastic recombination. Reduced Pgen has been associated with antigen-driven clonal convergence in viral infections such as COVID-19, where low Pgen, high-frequency clones reflect strong immune responses to specific viral epitopes (Schultheiß et al. 2020). Together, these observations support the interpretation that MS-restricted shared clones may arise under disease-specific selective pressures.

To identify potential antigenic factors in the B cell response associated with MS, we queried AgAbDB using HCDR3 sequences shared exclusively among MS patients, excluding clusters detected in HCs. The matched antibodies targeted both exogenous and endogenous antigens, consistent with the long-standing hypothesis that MS pathogenesis involves a complex interplay between infectious triggers and autoimmunity (Ascherio and Munger 2007). Among exogenous targets, we identified viral, bacterial, parasitic, and toxin-associated proteins. This is consistent with both experimental and epidemiological evidence of the association between viral agents, especially EBV, and MS (Bjornevik et al. 2022). However, the presence of multiple endogenous targets suggests the possibility of an extended spectrum of self-proteins in the context of MS-related autoimmunity. This is consistent with the phenomenon of epitope spreading, i.e., the diversification of epitope specificity from the initial immune response directed against a self or foreign protein to subdominant and/or cryptic epitopes on that protein or other proteins. Studies conducted on experimental models of MS show that epitope spreading plays a pathological role in the disease and that blocking this process inhibits ongoing clinical courses (Vanderlugt and Miller 2002).

Importantly, using Levenshtein distance thresholds (≤2 amino acid differences) allowed us to capture not only exact matches but also near-identical and similar sequences, potentially representing clonally related BCRs that may have diverged through somatic hypermutation and affinity maturation toward shared epitopes.

The identification of these convergent, putative antigen-specific signatures provides a foundation for future studies to better understand their role in MS pathogenesis.

This study has several limitations. The database is built entirely from previously published datasets that differ in sampling strategies, clinical inclusion criteria, and metadata richness. Although BCR sequence processing was standardized (Smakaj et al. 2020), substantial heterogeneity remains across studies in sequencing platforms, library preparation methods, primer sets, and cell-sorting workflows (Mhanna et al. 2024; Greiff et al. 2015; Miho et al. 2018; Stervbo et al. 2025). Metadata availability is inconsistent across datasets, limiting our ability to account for factors such as treatment history or disease duration. Because paired heavy- and light-chain information was not consistently available, our analyses were restricted to the heavy chain alone, reducing the precision of clonotype reconstruction and preventing a definitive inference of antigen specificity. The procedure used to match MS-specific BCR sequences to known antigen specificities relied exclusively on the HCDR3 amino acid region. While HCDR3 often represents the primary determinant of antigen binding, this approach may be less accurate for short HCDR3s or in cases where specificity is influenced by regions outside the HCDR3, such as CDR1, CDR2, or framework regions (Xu and Davis 2000; Akbar et al. 2021). Finally, although this is not a limitation of the study design per se, the database is inherently incomplete because some relevant datasets are not publicly accessible due to GDPR and data-sharing restrictions.

## Material and Methods

### Systematic review

We conducted a systematic review following PRISMA guidelines (Page et al.,(Page et al. 2021) to identify and collect raw BCR sequencing data from MS patients.

The research question was framed according to the PICO framework:

- Population (P): patients diagnosed with MS.
- Intervention/exposure (I): BCR sequencing
- Comparison (C): healthy controls, when available.
- Outcomes (O): (i) compilation of a comprehensive MS BCR database including all available raw sequencing data, and (ii) identification of potential disease-related BCR signatures across different individuals and tissues.

From database inception through 14 July 2025, we searched PubMed and EMBASE using the query: (MS OR “multiple sclerosis”) AND (BCR OR “B cell receptor” OR “B-cell receptor” OR “BCR repertoire” OR “B cell receptor repertoire” OR BCR sequenc* OR “B cell receptor sequenc*” OR “immunoglobulin heavy chain”). We also queried the iReceptor Gateway, NCBI Sequence Read Archive (SRA), and VDJServer for publicly available MS-related BCR datasets. Records were screened and deduplicated using Rayyan (Ouzzani et al. 2016). A PRISMA flowchart (Fig. 1a) summarizes the numbers of records identified, screened, excluded and included. Inclusion criteria encompassed studies analyzing human MS BCR data with available raw sequences published after 2005, corresponding to the advent of next-generation sequencing technologies (Margulies et al. 2005); whereas studies using animal models, non-MS cohorts, or without accessible raw data were excluded.

### MS-BCR Database construction

Raw .fastq files were obtained from the SRA when available or requested directly from authors when necessary. All datasets were processed with MiXCR (v4.6.0), using presets selected according to the sequencing strategy: 10x Genomics (Ramesh et al. 2020; Agrafiotis et al. 2023), amplicon (bulk sequencing studies: (Ryback and Cowan 2025; Palanichamy et al. 2014; Laurent et al. 2023; Stern et al. 2014; Lomakin et al. 2022; Greenfield et al. 2019; Saldivar et al. 2022; Zvyagin, n.d.; Ruschil et al. 2023). Germline gene assignment and framework/CDR region annotation were performed using the IMGT reference database and standardized IMGT nomenclature implemented within MiXCR. Processed data were exported in AIRR-compliant format, and erroneously included TCR sequences were removed.

Data from the original studies were retained and complemented with a dataset-level sequence quality score derived from FastQC’s per-base sequence quality module summarizing average Phred scores across raw sequencing reads (“Babraham Bioinformatics - FastQC A Quality Control Tool for High Throughput Sequence Data,” n.d.).

Demographic information was integrated, retaining when available, the following fields: disease, age, sex, analyzed tissue. Finally, all datasets were integrated into a single unified database. A unique sequence_id was assigned to each entry across the entire dataset to ensure consistent indexing and traceability. The integrated database is distributed in a modular format to facilitate usability and scalability. The complete MS-BCR-DB is publicly available and stored at Zenodo (DOI: 10.5281/zenodo.18862069 ), and the documentation and instructions for use, are available via GitHub at: https://github.com/chiarball/MS_BCR.

## Data analysis

### Cohort selection and BCR repertoire analysis in MS

We focused our analysis on naïve MS patients and incorporated a peripheral-blood repertoire from HCs (Ghraichy et al. 2020). Exploratory assessments of overall data distribution, junction amino-acid length, and V-D-J gene usage were performed in R (v4.5.0) and were graphically represented with the ggplot2 package (v3.5.2). Statistical comparisons of junction lengths and gene-usage frequencies were conducted in R using the Wilcoxon test or the Kruskal-Wallis test, as appropriate (p < 0.05, adjusted for multiple comparisons using the Benjamini-Hochberg). We defined clonotype as a unique combination of V and J genes and HCDR3 amino acid sequence. Clonal clustering was executed for each patient using ATrieGC software (v0.0.4) (Spisak et al. 2022), grouping sequences with identical V and J gene assignments and linked one to each other with a maximum amino-acid HCDR3 Hamming distance = 1. Cluster abundance was quantified as the number of sequences assigned to each cluster; clonal cluster repertoire proportions were then visualized with ggplot2.

### BCR clonal clustering across patients and network representation

We conducted a second clustering analysis using ATrieGC by pooling all sequences from patients and HCs. In contrast to the previous analysis, clustering was performed across all subjects rather than within each subject. Sequences were grouped by identical V and J gene assignments, and a link was assigned between two sequences when their amino-acid HCDR3 regions differed by a maximum Hamming distance of 1 (Spisak et al. 2022). Clusters containing at least one sequence from HCs were then excluded to avoid including public clonotypes that were present in the healthy repertoire. Only clusters composed exclusively of patient-derived sequences were retained. The representation of clusters as networks (Fig. 2B) was done in Python (v3.12.7) via NetworkX (v3.3). To select clusters for network representation, we developed a composite scoring system designed to emphasize sharing across patients and tissues while accounting for cluster size and distributional balance. For each cluster, inter-patient sharedness and inter-tissue dissemination were defined as the proportion of unique patients and tissues represented, respectively. The inter-patient sharedness component was weighted by Pielou’s evenness (Pielou 1966), computed from the distribution of sequences across patients. Cluster size was defined as the number of unique sequences per cluster per patient, summed across patients and normalized to the maximum cluster size in the dataset. The final score was calculated as a product of these components (*sharedness * sharedness evenness * dissemination * cluster size*).

### Lineage reconstruction, phylogenetic analysis and Pgen estimation

Clonal lineages were inferred with HILARy (v1.2.3), a high-precision method that integrates probabilistic models of V(D)J recombination and selection with clustering of shared somatic hypermutations to identify B-cell clonal families (Spisak et al. 2022). RAxML (v.8.2.13) (Stamatakis 2014) was used to reconstruct phylogenetic trees rooted on the inferred germline sequence for each lineage; lineage trees were visualized in iTOL (Letunic and Bork 2024).

To quantify the likelihood of observing MS shared sequences (present in >1 patient) solely through stochastic recombination, we computed their generation probability (Pgen), defined as the probability of a given sequence arising through V(D)J recombination, using OLGA (v1.2.4) (Sethna et al. 2019).

### Matching MS-BCR-DB sequences to the harmonized AgAb database

To investigate whether MS-BCR sequences match those of known antigen specificity, we queried the Antigen-specific Antibody Database (Czerwiński et al. 2025),(Czerwiński et al. 2025); https://naturalantibody.com/agab/), downloaded on the 5th of August 2025.

To harmonize antigen annotation in the AgAb database, antibody sequences were grouped by identical HCDR3 amino acid sequence, IGHV gene, and mutation status (≥98% germline identity classified as unmutated). Antigen annotations from patent sources were normalized by lowercasing and removal of parenthetical content and non-informative punctuation. Annotation consistency within each HCDR3 group was assessed using hierarchical clustering of antigen strings with the Jaro-Winkler distance (cut height h ≤ 0.20, (Winkler 1990); cluster-level semantic purity, defined as the proportion of the most frequent annotation within each group, was then used as a quantitative measure of annotation consistency. Groups annotated with one or two antigens, or high semantic purity (≥0.7) were retained. Groups with average purity between 0.5 and 0.7 were retained only if they had limited annotation counts (≤10). Groups that did not satisfy the aforementioned criteria were excluded from further analysis. For retained groups, heterogeneous antigen annotations were consolidated into a single representative antigen label using a bag-of-words approach. The resulting naming variants were harmonized with the rest of the database annotations via a token-based Jaccard similarity and graph clustering. To prevent confounding due to public or shared clonotypes present in HCs, only clusters shared among MS patients were retained for antigen-association analysis. Pairwise comparisons between filtered MS HCDR3 sequences and AgAbDb sequences were performed using the Levenshtein distance, retaining matches with a distance ≤ 2. For each AgAbDb HCDR3, the minimum observed distance to any MS sequence was used to classify evidence levels into three groups: exact matches (distance = 0), near-identical (distance = 1), and similar (distance = 2). Antigen enrichment analysis was then performed in Python (v3.12.7). Fisher’s exact tests compared the proportion of MS-matched sequences associated with each antigen to their overall frequency in the AgAbDb background (Amoriello et al. 2021). P-values were corrected for multiple testing using the Benjamini-Hochberg false discovery rate (FDR) procedure, and standardized Pearson residuals were computed to quantify the strength of enrichment for each antigen. Visualization with boxplots of the significant match was made with the ggplot2 package (v3.5.2) in R (v4.5.0). Code and documentation associated with this study are publicly accessible via GitHub at: https://github.com/chiarball/MS_BCR/tree/main.

## Authors’ Contributions

Ch.B.: conceptualization, data analysis, data interpretation, figures, and writing; N.C.: conceptualization, data analysis, data interpretation, figures, and writing; M.F.A.: interpretation of results and manuscript revision; K.L.Q.: data preprocessing with MiXCR and analysis support; N.R.: database curation and quality control; L.W.: data analysis and preparation of supplementary materials; A.L.: conceptualization and manuscript revision; E.P.: manuscript revision; M.P.A.: manuscript revision; Cl.B.: conceptualization and manuscript revision; V.G.: study conceptualization, data interpretation, and writing.

## Bibliography

Afzali, Ali Maisam, and Thomas Korn. 2025. “The Role of the Adaptive Immune System in the Initiation and Persistence of Multiple Sclerosis.” Seminars in Immunology 78 (101947): 101947.

Agrafiotis, Andreas, Raphael Dizerens, Ilena Vincenti, et al. 2023. “Persistent Virus-Specific and Clonally Expanded Antibody-Secreting Cells Respond to Induced Self-Antigen in the CNS.” Acta Neuropathologica 145 (3): 335–355.

Akbar, Rahmad, Philippe A. Robert, Milena Pavlović, et al. 2021. “A Compact Vocabulary of Paratope-Epitope Interactions Enables Predictability of Antibody-Antigen Binding.” Cell Reports 34 (11): 108856.

Al Kindi, Mahmood A., Tim K. Chataway, George A. Gilada, et al. 2015. “Serum SmD Autoantibody Proteomes Are Clonally Restricted and Share Variable-Region Peptides.” Journal of Autoimmunity 57 (February): 77–81.

Amoriello, Roberta, Maria Chernigovskaya, Victor Greiff, et al. 2021. “TCR Repertoire Diversity in Multiple Sclerosis: High-Dimensional Bioinformatics Analysis of Sequences from Brain, Cerebrospinal Fluid and Peripheral Blood.” EBioMedicine 68 (103429): 103429.

Ascherio, Alberto, and Kassandra L. Munger. 2007. “Environmental Risk Factors for Multiple Sclerosis. Part I: The Role of Infection.” Annals of Neurology 61 (4): 288–299.

Avnir, Yuval, Aimee S. Tallarico, Quan Zhu, et al. 2014. “Molecular Signatures of Hemagglutinin Stem-Directed Heterosubtypic Human Neutralizing Antibodies against Influenza A Viruses.” PLoS Pathogens 10 (5): e1004103.

Avnir, Yuval, Corey T. Watson, Jacob Glanville, et al. 2016. “IGHV1-69 Polymorphism Modulates Anti-Influenza Antibody Repertoires, Correlates with IGHV Utilization Shifts and Varies by Ethnicity.” Scientific Reports 6 (1): 20842.

“Babraham Bioinformatics - FastQC A Quality Control Tool for High Throughput Sequence Data.” n.d. Accessed February 20, 2026. https://www.bioinformatics.babraham.ac.uk/projects/fastqc/.

Bankoti, Jaishree, Leonard Apeltsin, Stephen L. Hauser, et al. 2014. “In Multiple Sclerosis, Oligoclonal Bands Connect to Peripheral B-Cell Responses: OCB Connection to Periphery.” Annals of Neurology 75 (2): 266–276.

Baranzini, S. E., M. C. Jeong, C. Butunoi, R. S. Murray, C. C. Bernard, and J. R. Oksenberg. 1999. “B Cell Repertoire Diversity and Clonal Expansion in Multiple Sclerosis Brain Lesions.” The Journal of Immunology 163 (9): 5133–5144.

Bashford-Rogers, Rachael J. M., Kenneth G. C. Smith, and David C. Thomas. 2018. “Antibody Repertoire Analysis in Polygenic Autoimmune Diseases.” Immunology 155 (1): 3–17.

Bjornevik, Kjetil, Marianna Cortese, Brian C. Healy, et al. 2022. “Longitudinal Analysis Reveals High Prevalence of Epstein-Barr Virus Associated with Multiple Sclerosis.” Science (New York, N.Y.) 375 (6578): 296–301.

Blazso, Peter, Krisztian Csomos, Christopher M. Tipton, Boglarka Ujhazi, and Jolan E. Walter. 2022. “Lineage Reconstruction of in Vitro Identified Antigen-Specific Autoreactive B Cells from Adaptive Immune Receptor Repertoires.” International Journal of Molecular Sciences 24 (1): 225.

Büdingen, H-Christian von, Tracy C. Kuo, Marina Sirota, et al. 2012. “B Cell Exchange across the Blood-Brain Barrier in Multiple Sclerosis.” The Journal of Clinical Investigation 122 (12): 4533–4543.

Centonze, D., L. Muzio, S. Rossi, R. Furlan, G. Bernardi, and G. Martino. 2010. “The Link between Inflammation, Synaptic Transmission and Neurodegeneration in Multiple Sclerosis.” Cell Death and Differentiation 17 (7): 1083–1091.

Chatzidimitriou, Anastasia, Eva Minga, Thomas Chatzikonstantinou, Carol Moreno, Kostas Stamatopoulos, and Paolo Ghia. 2020. “Challenges and Solutions for Collecting and Analyzing Real World Data: The Eric CLL Database as an Illustrative Example.” HemaSphere 4 (5): e425.

Cremin, Conor John, Sabyasachi Dash, and Xiaofeng Huang. 2022. “Big Data: Historic Advances and Emerging Trends in Biomedical Research.” Current Research in Biotechnology 4: 138–151.

Czerwiński, Arkadiusz, Paweł Dudzic, Konrad Wójtowicz, et al. 2025. “ASD: Antigen-Specific Antibody Database.” In Bioinformatics, No. Biorxiv;2025.11.24.690097v1. BioRxiv, November 26. https://www.biorxiv.org/content/10.1101/2025.11.24.690097v1.

Dendrou, Calliope A., Lars Fugger, and Manuel A. Friese. 2015. “Immunopathology of Multiple Sclerosis.” Nature Reviews. Immunology 15 (9): 545–558.

Di Sabatino, Elena, Diana Ferraro, Lorenzo Gaetani, Edoardo Emiliano, Lucilla Parnetti, and Massimiliano Di Filippo. 2025. “CSF Biomarkers of B-Cell Activation in Multiple Sclerosis: A Clinical Perspective.” Journal of Neurology 272 (3): 211.

Ghraichy, Marie, Jacob D. Galson, Aleksandr Kovaltsuk, et al. 2020. “Maturation of the Human Immunoglobulin Heavy Chain Repertoire with Age.” Frontiers in Immunology 11 (August): 1734.

Greenfield, Ariele L., Ravi Dandekar, Akshaya Ramesh, et al. 2019. “Longitudinally Persistent Cerebrospinal Fluid B Cells Can Resist Treatment in Multiple Sclerosis.” JCI Insight 4 (6). 10.1172/jci.insight.126599.

Greiff, Victor, Enkelejda Miho, Ulrike Menzel, and Sai T. Reddy. 2015. “Bioinformatic and Statistical Analysis of Adaptive Immune Repertoires.” Trends in Immunology 36 (11): 738–749.

Guo, Huadong. 2015. “Big Data for Scientific Research and Discovery.” International Journal of Digital Earth 8 (1): 1–2.

Hauser, Stephen L. 2015. “The Charcot Lecture | Beating MS: A Story of B Cells, with Twists and Turns.” Multiple Sclerosis (Houndmills, Basingstoke, England) 21 (1): 8–21.

Horns, Felix, Christopher Vollmers, Derek Croote, et al. 2016. “Lineage Tracing of Human B Cells Reveals the in Vivo Landscape of Human Antibody Class Switching.” eLife 5 (August). 10.7554/eLife.16578.

Ishigaki, Kazuyoshi, Kaitlyn A. Lagattuta, Yang Luo, Eddie A. James, Jane H. Buckner, and Soumya Raychaudhuri. 2022. “HLA Autoimmune Risk Alleles Restrict the Hypervariable Region of T Cell Receptors.” Nature Genetics 54 (4): 393–402.

Jelcic, Ivan, Reza Naghavian, Imran Fanaswala, et al. 2025. “T-Bet+ CXCR3+ B Cells Drive Hyperreactive B-T Cell Interactions in Multiple Sclerosis.” Cell Reports. Medicine 6 (3): 102027.

John, Gareth R., Sai Latha Shankar, Bridget Shafit-Zagardo, et al. 2002. “Multiple Sclerosis: Re-Expression of a Developmental Pathway That Restricts Oligodendrocyte Maturation.” Nature Medicine 8 (10): 1115–1121.

Kurowska, Z., P. Brundin, M. E. Schwab, and J-Y Li. 2014. “Intracellular Nogo-A Facilitates Initiation of Neurite Formation in Mouse Midbrain Neurons in Vitro.” Neuroscience 256 (January): 456–466.

Lanz, Tobias V., R. Camille Brewer, Peggy P. Ho, et al. 2022. “Clonally Expanded B Cells in Multiple Sclerosis Bind EBV EBNA1 and GlialCAM.” Nature 603 (7900): 321–327.

Laurent, Sarah A., Nicolas B. Strauli, Erica L. Eggers, et al. 2023. “Effect of Ocrelizumab on B- and T-Cell Receptor Repertoire Diversity in Patients with Relapsing Multiple Sclerosis from the Randomized Phase III OPERA Trial.” Neurology(R) Neuroimmunology & Neuroinflammation 10 (4). 10.1212/NXI.0000000000200118.

Letunic, Ivica, and Peer Bork. 2024. “Interactive Tree of Life (iTOL) v6: Recent Updates to the Phylogenetic Tree Display and Annotation Tool.” Nucleic Acids Research 52 (W1): W78–W82.

Lindeman, Ida, Justyna Polak, Shuo-Wang Qiao, et al. 2022. “Stereotyped B-Cell Responses Are Linked to IgG Constant Region Polymorphisms in Multiple Sclerosis.” European Journal of Immunology 52 (4): 550–565.

Li, Rui, Kristina R. Patterson, and Amit Bar-Or. 2018. “Reassessing B Cell Contributions in Multiple Sclerosis.” Nature Immunology 19 (7): 696–707.

Lisak, Robert P., Joyce A. Benjamins, Liljana Nedelkoska, et al. 2012. “Secretory Products of Multiple Sclerosis B Cells Are Cytotoxic to Oligodendroglia in Vitro.” Journal of Neuroimmunology 246 (1-2): 85–95.

Lisak, Robert P., Liljana Nedelkoska, Joyce A. Benjamins, et al. 2017. “B Cells from Patients with Multiple Sclerosis Induce Cell Death via Apoptosis in Neurons in Vitro.” Journal of Neuroimmunology 309 (August): 88–99.

Liu, Hongmei, Wenjing Pan, Congli Tang, et al. 2021. “The Methods and Advances of Adaptive Immune Receptors Repertoire Sequencing.” Theranostics 11 (18): 8945–8963.

Lomakin, Y. A., L. A. Ovchinnikova, M. N. Zakharova, et al. 2022. “Multiple Sclerosis Is Associated with Immunoglobulin Germline Gene Variation of Transitional B Cells.” Acta Naturae 14 (4): 84–93.

Lovato, Laura, Simon N. Willis, Scott J. Rodig, et al. 2011. “Related B Cell Clones Populate the Meninges and Parenchyma of Patients with Multiple Sclerosis.” Brain: A Journal of Neurology 134 (Pt 2): 534–541.

Luchicchi, Antonio, Paolo Preziosa, and Bert ‘t Hart. 2021. “Editorial: ‘Inside-out’ vs ‘Outside-in’ Paradigms in Multiple Sclerosis Etiopathogenesis.” Frontiers in Cellular Neuroscience 15 (March): 666529.

Magliozzi, Roberta, Owain Howell, Abhilash Vora, et al. 2007. “Meningeal B-Cell Follicles in Secondary Progressive Multiple Sclerosis Associate with Early Onset of Disease and Severe Cortical Pathology.” Brain: A Journal of Neurology 130 (Pt 4): 1089–1104.

Marti, Zoe, Josefine Ruder, Olivia G. Thomas, et al. 2024. “Enhanced and Cross-Reactive in Vitro Memory B Cell Response against Epstein-Barr Virus Nuclear Antigen 1 in Multiple Sclerosis.” Frontiers in Immunology 15 (August): 1334720.

McArthur, Simon, Rodrigo Azevedo Loiola, Elisa Maggioli, Mariella Errede, Daniela Virgintino, and Egle Solito. 2016. “The Restorative Role of Annexin A1 at the Blood-Brain Barrier.” Fluids and Barriers of the CNS 13 (1): 17.

Mhanna, Vanessa, Habib Bashour, Khang Lê Quý, et al. 2024. “Adaptive Immune Receptor Repertoire Analysis.” Nature Reviews Methods Primers 4 (1). 10.1038/s43586-023-00284-1.

Miho, Enkelejda, Alexander Yermanos, Cédric R. Weber, Christoph T. Berger, Sai T. Reddy, and Victor Greiff. 2018. “Computational Strategies for Dissecting the High-Dimensional Complexity of Adaptive Immune Repertoires.” Frontiers in Immunology 9 (February): 224.

Mora, Pierre, and Candice Chapouly. 2023. “Astrogliosis in Multiple Sclerosis and Neuro-Inflammation: What Role for the Notch Pathway?” Frontiers in Immunology 14 (October): 1254586.

Oh, Jiwon, Angela Vidal-Jordana, and Xavier Montalban. 2018. “Multiple Sclerosis: Clinical Aspects.” Current Opinion in Neurology 31 (6): 752–759.

Ouzzani, Mourad, Hossam Hammady, Zbys Fedorowicz, and Ahmed Elmagarmid. 2016. “Rayyan-a Web and Mobile App for Systematic Reviews.” Systematic Reviews 5 (1): 210.

Owens, Gregory P., Jeffrey L. Bennett, Donald H. Gilden, and Mark P. Burgoon. 2006. “The B Cell Response in Multiple Sclerosis.” Neurological Research 28 (3): 236–244.

Owens, Gregory P., Kimberly M. Winges, Alanna M. Ritchie, et al. 2007. “VH4 Gene Segments Dominate the Intrathecal Humoral Immune Response in Multiple Sclerosis.” The Journal of Immunology 179 (9): 6343–6351.

Page, Matthew J., Joanne E. McKenzie, Patrick M. Bossuyt, et al. 2021. “The PRISMA 2020 Statement: An Updated Guideline for Reporting Systematic Reviews.” BMJ (Clinical Research Ed.) 372 (March): n71.

Palanichamy, Arumugam, Leonard Apeltsin, Tracy C. Kuo, et al. 2014. “Immunoglobulin Class-Switched B Cells Form an Active Immune Axis between CNS and Periphery in Multiple Sclerosis.” Science Translational Medicine 6 (248): 248ra106.

Peng, Ge, Carlo Lacagnina, Robert R. Downs, et al. 2022. “Global Community Guidelines for Documenting, Sharing, and Reusing Quality Information of Individual Digital Datasets.” Data Science Journal 21 (March). 10.5334/dsj-2022-008.

Pérez-Saldívar, Miriam, Yusuke Nakamura, Kazuma Kiyotani, et al. 2024. “Comparative Analysis of the B Cell Receptor Repertoire during Relapse and Remission in Patients with Multiple Sclerosis.” Clinical Immunology (Orlando, Fla.) 269 (110398): 110398.

Pielou, E. C. 1966. “The Measurement of Diversity in Different Types of Biological Collections.” Journal of Theoretical Biology 13 (December): 131–144.

Qin, Y., P. Duquette, Y. Zhang, P. Talbot, R. Poole, and J. Antel. 1998. “Clonal Expansion and Somatic Hypermutation of V(H) Genes of B Cells from Cerebrospinal Fluid in Multiple Sclerosis.” The Journal of Clinical Investigation 102 (5): 1045–1050.

Ramesh, Akshaya, Ryan D. Schubert, Ariele L. Greenfield, et al. 2020. “A Pathogenic and Clonally Expanded B Cell Transcriptome in Active Multiple Sclerosis.” Proceedings of the National Academy of Sciences of the United States of America 117 (37): 22932–22943.

Rashidbenam, Zahra, Ezgi Ozturk, Maurice Pagnin, Paschalis Theotokis, Nikolaos Grigoriadis, and Steven Petratos. 2023. “How Does Nogo Receptor Influence Demyelination and Remyelination in the Context of Multiple Sclerosis?” Frontiers in Cellular Neuroscience 17 (June): 1197492.

Raybould, Matthew I. J., Aleksandr Kovaltsuk, Claire Marks, and Charlotte M. Deane. 2021. “CoV-AbDab: The Coronavirus Antibody Database.” Bioinformatics 37 (5): 734–735.

Rivas, Jacqueline R., Sara J. Ireland, Rati Chkheidze, et al. 2017. “Peripheral VH4+ Plasmablasts Demonstrate Autoreactive B Cell Expansion toward Brain Antigens in Early Multiple Sclerosis Patients.” Acta Neuropathologica 133 (1): 43–60.

Rossi, Silvia, Valeria Studer, Alessandro Moscatelli, et al. 2013. “Opposite Roles of NMDA Receptors in Relapsing and Primary Progressive Multiple Sclerosis.” PloS One 8 (6): e67357.

Ruschil, Christoph, Gisela Gabernet, Constanze Louisa Kemmerer, et al. 2023. “Cladribine Treatment Specifically Affects Peripheral Blood Memory B Cell Clones and Clonal Expansion in Multiple Sclerosis Patients.” Frontiers in Immunology 14 (March): 1133967.

Ryback, Audrey A., and Graeme J. M. Cowan. 2025. “Deep Sequencing of BCR Heavy Chain Repertoires in Myalgic Encephalomyelitis/chronic Fatigue Syndrome.” Frontiers in Immunology 16 (February): 1489312.

Saldivar, Perez, M. P. Ordoñez, G. Sotelo, J. Martínez Palomo Flores, and J. Espinosa. 2022. Comparative Analysis B Cell Receptor Repertoire Patients Multiple Sclerosis during Relapse Remission.

Satoh, Jun-Ichi, Hiroyuki Onoue, Kunimasa Arima, and Takashi Yamamura. 2005. “Nogo-A and Nogo Receptor Expression in Demyelinating Lesions of Multiple Sclerosis.” Journal of Neuropathology and Experimental Neurology 64 (2): 129–138.

Schultheiß, Christoph, Lisa Paschold, Donjete Simnica, et al. 2020. “Next-Generation Sequencing of T and B Cell Receptor Repertoires from COVID-19 Patients Showed Signatures Associated with Severity of Disease.” Immunity 53 (2): 442–455.e4.

Serafini, Barbara, Barbara Rosicarelli, Roberta Magliozzi, Egidio Stigliano, and Francesca Aloisi. 2004. “Detection of Ectopic B-Cell Follicles with Germinal Centers in the Meninges of Patients with Secondary Progressive Multiple Sclerosis.” Brain Pathology (Zurich, Switzerland) 14 (2): 164–174.

Sethna, Zachary, Yuval Elhanati, Curtis G. Callan, Aleksandra M. Walczak, and Thierry Mora. 2019. “OLGA: Fast Computation of Generation Probabilities of B- and T-Cell Receptor Amino Acid Sequences and Motifs.” Bioinformatics (Oxford, England) 35 (17): 2974–2981.

Shi, Bin, Xiaoheng Dong, Qingqing Ma, et al. 2020. “The Usage of Human IGHJ Genes Follows a Particular Non-Random Selection: The Recombination Signal Sequence May Affect the Usage of Human IGHJ Genes.” Frontiers in Genetics 11 (524413): 524413.

Shiroishi, Mitsunori. 2021. “Structural Basis of a Conventional Recognition Mode of IGHV1-69 Rheumatoid Factors.” Advances in Experimental Medicine and Biology 21: 171–182.

Singh, Vaibhav, Marcel P. Stoop, Christoph Stingl, et al. 2013. “Cerebrospinal-Fluid-Derived Immunoglobulin G of Different Multiple Sclerosis Patients Shares Mutated Sequences in Complementarity Determining Regions.” Molecular & Cellular Proteomics: MCP 12 (12): 3924–3934.

Slatko, Barton E., Andrew F. Gardner, and Frederick M. Ausubel. 2018. “Overview of next-Generation Sequencing Technologies: Overview of next-Generation Sequencing.” Et Al [Current Protocols in Molecular Biology] 122 (1): e59.

Smakaj, Erand, Lmar Babrak, Mats Ohlin, et al. 2020. “Benchmarking Immunoinformatic Tools for the Analysis of Antibody Repertoire Sequences.” Bioinformatics (Oxford, England) 36 (6): 1731–1739.

Spender, Lindsay C., Georgina H. Cornish, Alexandra Sullivan, and Paul J. Farrell. 2002. “Expression of Transcription Factor AML-2 (RUNX3, CBF(alpha)-3) Is Induced by Epstein-Barr Virus EBNA-2 and Correlates with the B-Cell Activation Phenotype.” Journal of Virology 76 (10): 4919–4927.

Spisak, Natanael, Thomas Dupic, Thierry Mora, and Aleksandra M. Walczak. 2022. “Combining Mutation and Recombination Statistics to Infer Clonal Families in Antibody Repertoires.” In bioRxiv. December 22. arXiv. 10.1101/2022.12.22.521661.

Stamatakis, Alexandros. 2014. “RAxML Version 8: A Tool for Phylogenetic Analysis and Post-Analysis of Large Phylogenies.” Bioinformatics 30 (9): 1312–1313.

Steiman-Shimony, Avital, Hanna Edelman, Anat Hutzler, et al. 2006. “Lineage Tree Analysis of Immunoglobulin Variable-Region Gene Mutations in Autoimmune Diseases: Chronic Activation, Normal Selection.” Cellular Immunology 244 (2): 130–136.

Stern, Joel N. H., Gur Yaari, Jason A. Vander Heiden, et al. 2014. “B Cells Populating the Multiple Sclerosis Brain Mature in the Draining Cervical Lymph Nodes.” Science Translational Medicine 6 (248): 248ra107.

Stervbo, Ulrik, Paraskevas Filippidis, Felix Breden, et al. 2025. “Challenges and Future Directions of AIRR-Seq-Based Diagnostics.” Immunoinformatics (Amsterdam, Netherlands) 19 (100056): 100056.

Thomas, Olivia G., Mattias Bronge, Katarina Tengvall, et al. 2023. “Cross-Reactive EBNA1 Immunity Targets Alpha-Crystallin B and Is Associated with Multiple Sclerosis.” Science Advances 9 (20): eadg3032.

Vanderlugt, Carol L., and Stephen D. Miller. 2002. “Epitope Spreading in Immune-Mediated Diseases: Implications for Immunotherapy.” Nature Reviews Immunology 2 (2): 85–95.

Watson, C. T., and F. Breden. 2012. “The Immunoglobulin Heavy Chain Locus: Genetic Variation, Missing Data, and Implications for Human Disease.” Genes and Immunity 13 (5): 363–373.

Winkler, William E. 1990. String Comparator Metrics and Enhanced Decision Rules in the Fellegi-Sunter Model of Record Linkage. https://eric.ed.gov/?id=ED325505.

Xu, J. L., and M. M. Davis. 2000. “Diversity in the CDR3 Region of V(H) Is Sufficient for Most Antibody Specificities.” Immunity 13 (1): 37–45.

Younis, Shady, Sajede Rasouli, Jacob W. Loeffler, et al. 2026. “EBV Reprograms Autoreactive Anti-CNS B Cells as Antigen Presenting Cells in Multiple Sclerosis.” In bioRxiv. BioRxiv, February 12. 10.64898/2026.02.11.701910.

Zvyagin. n.d. “Comparative Analysis B Cell Receptor Repertoires Revealed Delay CD24hiCD38hi Transitional B Lymphocyte Maturation Multiple Sclerosis Patients.” Preprint.

