## Supplementary material for "MS-BCR-DB: an integrated BCR repertoire database to mine humoral multiple sclerosis signatures": supp_fig_and_table captions.pdf

**Supplementary Fig. 1** HCDR3 junction amino acid length distribution in MS patients across tissues and anatomical compartments **A** Line plots show the normalized distribution of junction amino acid lengths across individual tissues. **B** Box plots show junction amino acid length distributions in intracranial versus extracranial tissues. Intracranial samples include CSF and brain-derived tissues (pia mater, choroid plexus, brain lesion), while extracranial samples include blood and CLN.
Statistical significance was assessed using Kruskal-Wallis tests for comparisons across all tissues and Wilcoxon rank-sum tests for intracranial versus extracranial comparisons, with Benjamini-Hochberg false discovery rate (FDR) correction for multiple testing ($p < 0.05$ *, $p < 0.01$ **, $p < 0.001$ ***). The junction region was defined according to the IMGT criteria, including the conserved Cysteine (C) and Tryptophan (W) residues, consistent with the AIRR Community data format specifications
Abbreviations: CLN, cervical lymph nodes; CSF, cerebrospinal fluid.

**Supplementary Fig. 2** Distribution of V(D)J gene usage across extracranial and intracranial tissues in MS patients. Box plots show per-subject relative gene usage frequencies, computed within each anatomical compartment and normalized by total repertoire size. Intracranial samples include CSF and brain-derived tissues (pia mater, choroid plexus, brain lesion), while extracranial samples include blood and CLN. **A** Individual V genes. **B** V gene families. **C** Individual D genes. **D** Individual J genes. Statistical significance was assessed per-gene (or gene-family) using Wilcoxon rank-sum tests with Benjamini-Hochberg FDR correction for multiple comparisons ($p < 0.05$ *, $p < 0.01$ **, $p < 0.001$ ***).

**Supplementary Fig. 3** Distribution of V(D)J gene usage and HCDR3 junction length in MS patients versus HCs. Box plots show per-subject relative frequencies, computed within each repertoire and normalized by total repertoire size. **A** V genes. **B** D genes. **C** J genes. **D** junction amino acid length distribution. Statistical significance was assessed per-gene (or per-length) using Wilcoxon rank-sum tests with Benjamini-Hochberg false discovery rate (FDR) correction for multiple comparisons ($p < 0.05$ *, $p < 0.01$ **, $p < 0.001$ ***). The junction region was defined according to the IMGT criteria, including the conserved Cysteine (C) and Tryptophan (W) residues, consistent with the AIRR Community data format specifications.

**Supplementary Fig. 4** Distribution of the probability of generation (Pgen, log10 scale) of BCR sequences shared by at least two subjects, stratified by their presence exclusively in MS clusters or shared between MS patients and healthy controls (HCs). Sequences shared with HCs show higher Pgen values, consistent with a higher probability of generation, whereas sequences detected only in MS clusters show lower Pgen values, suggesting a more restricted or disease-associated origin.

**Supplementary Table 1** Overview of BCR datasets included in this study. This supplementary table provides a comprehensive overview of all BCR sequencing datasets analyzed in this study and is organized into multiple sheets. The sheet S1_*General_information* summarizes study characteristics, including cohort composition and clinical context, biological material and B cell subsets profiled, key experimental features of BCR sequencing, and data availability. The sheet S2_*Clinical_metadata_all* compiles all available clinical metadata across the different studies in a harmonized format. The remaining sheets (*S3-S12)* report the detailed clinical information available for each study. Abbreviations: DMT, disease-modifying therapy (defined as any form of immunomodulatory or immunosuppressive treatment); PBMCs, peripheral blood mononuclear cells; CSF, cerebrospinal fluid; CNS, central nervous system; CLN, cervical lymph nodes; VH, variable heavy chain; CH, constant heavy chain; VL, variable light chain; CL, constant light chain.

**Supplementary Table 2** Complete list of Pgen calculations and metadata for all MS-associated clusters. Clonotypes were defined as immunoglobulin heavy-chain sequences sharing identical IGHV gene, IGHJ gene, and HCDR3 amino acid sequence. Clonotype clusters were generated using single-linkage clustering. Clonotypes were assigned to the same cluster if they shared identical IGHV and IGHJ gene usage and were connected by at least one pairwise link corresponding to a Hamming distance of 1 within the HCDR3 amino acid sequence.The column *cluster_h1* reports the unique identifier of the clonotype cluster. *Pgen* indicates the generation probability of the HCDR3 sequence as estimated by OLGA probabilistic recombination modeling. *In_top_cluster* specifies whether the corresponding cluster is included among the top 100 clusters displayed in Figure 2B. Additional columns report the tissue of origin, subject identifier, and study cohort.

**Supplementary Table 3** FASTQ sequencing quality metrics. Summary of FASTQ sequencing quality metrics across the analyzed studies based on Phred quality scores. Quality metrics were calculated for individual sequencing runs (SRR entries) and then summarized at the study level. The table reports the number of samples per study (*sample_count*) and summary statistics of sequencing depth (*total_sequences*), GC content (*gc_percent*), mean Phred quality scores (*mean_quality_score*), and the proportion of bases with quality ≥Q30 (*percent_bases_above_q30*), reported as minimum, maximum, median, mean, or standard deviation across samples. Additional columns report the number of samples flagged for adapter contamination (*adapter_contamination_flag_fail_count*) and the percentage of samples passing the overall FastQC quality assessment (*fastqc_overall_status_pass_percent*).
