## Supplementary material for "MS-BCR-DB: an integrated BCR repertoire database to mine humoral multiple sclerosis signatures": supplementary_figs_all.pdf

### Supplementary Figure 1

A Junction amino acid length distribution across tissues in MS

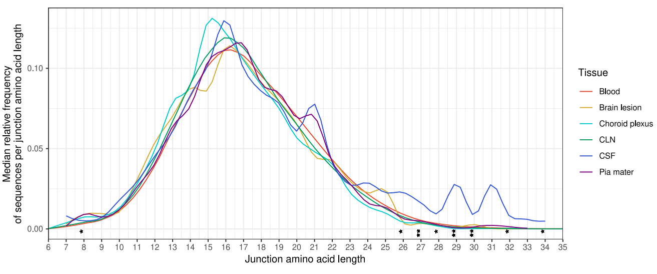


B Junction amino acid length in intra- and extra-cranial compartments

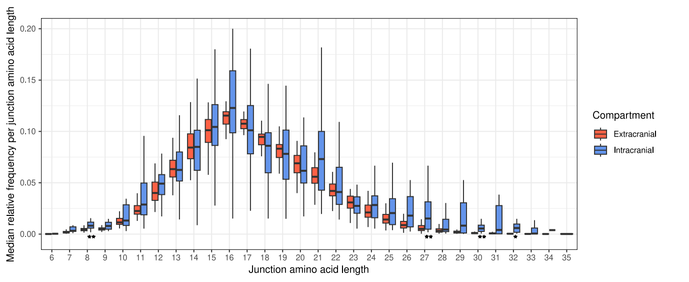


### Supplementary Figure 2

A IGHV gene usage across anatomical compartments in MS

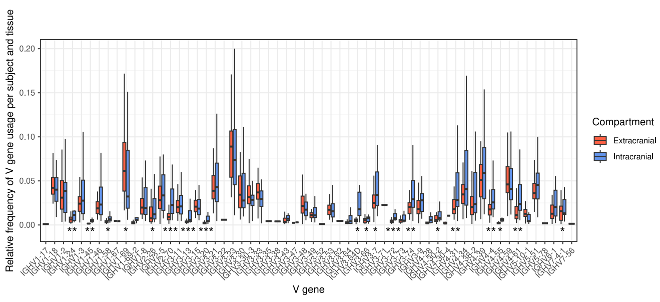


B IGHV family usage across anatomical compartments in MS

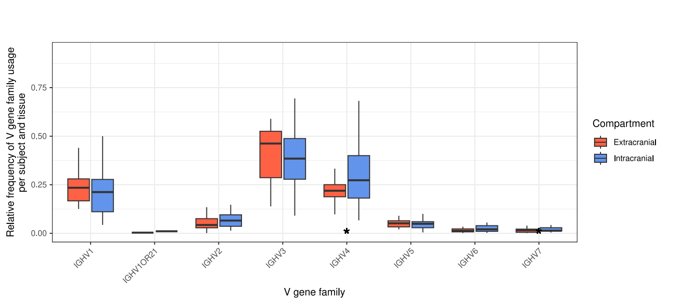


C IGHD gene usage across anatomical compartments in MS

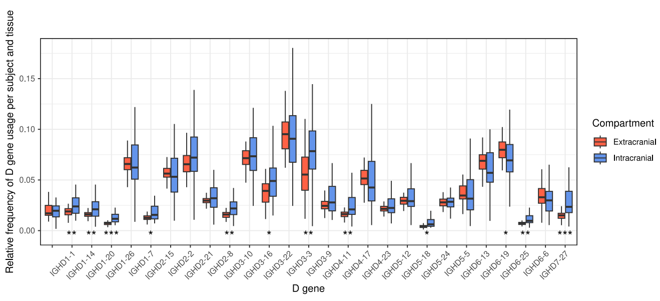


D IGHJ gene usage across anatomical compartments in MS

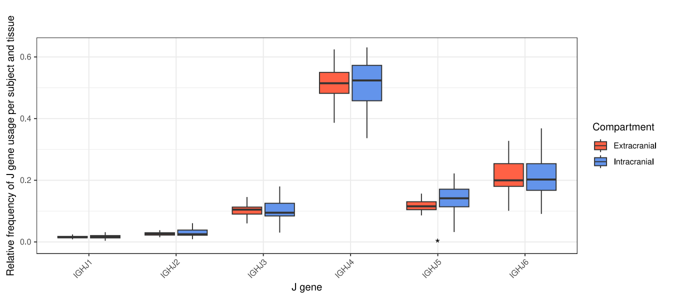


### Supplementary Figure 3

A IGHV gene usage in MS versus HCs

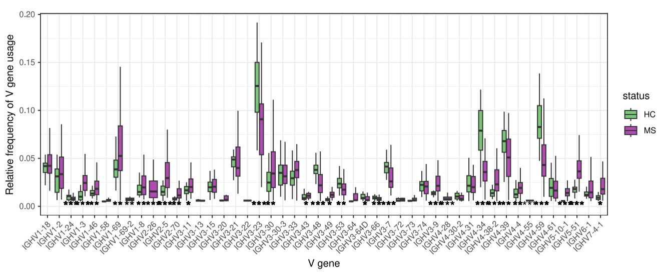


B IGHD gene usage in MS versus HCs

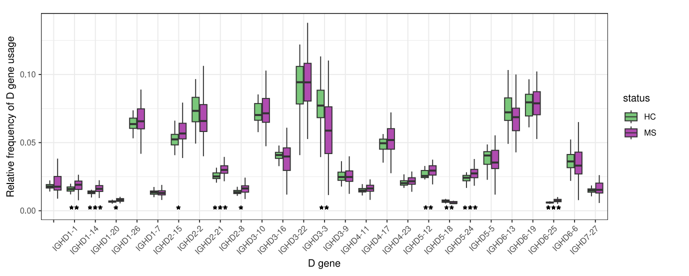


C IGHJ gene usage in MS versus HCs

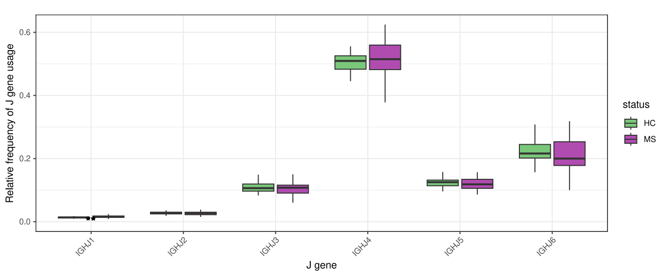


D Junction amino acid length in MS versus HCs

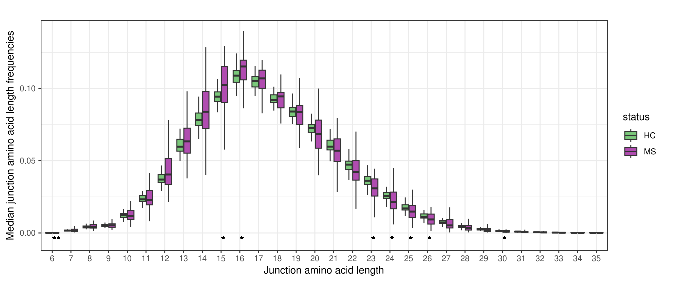


### Supplementary Figure 4

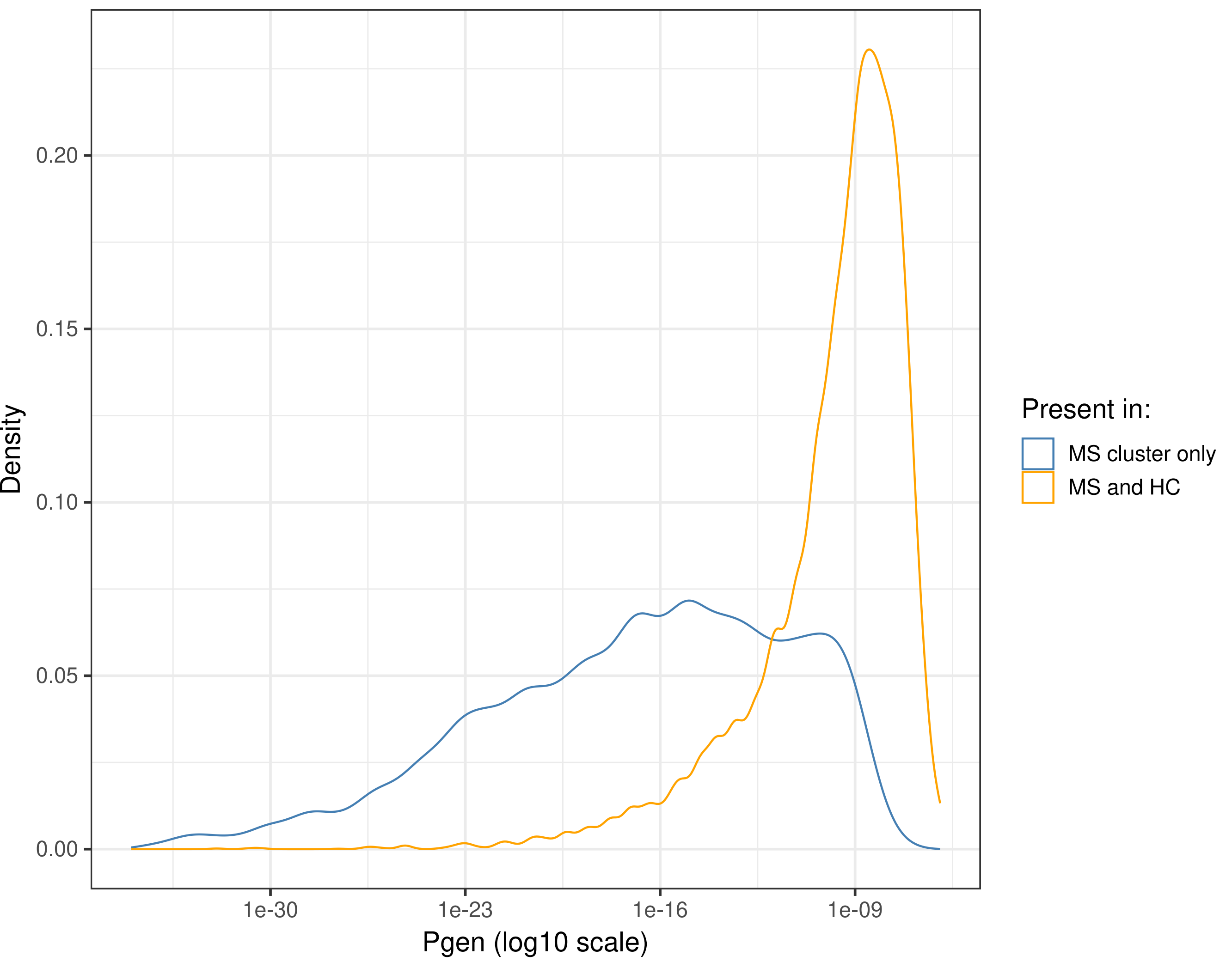
